# A single cell-based atlas of human microglial states reveals associations with neurological disorders and histopathological features of the aging brain

**DOI:** 10.1101/343780

**Authors:** Marta Olah, Vilas Menon, Naomi Habib, Mariko Taga, Christina Yung, Maria Cimpean, Anthony Khairalla, Danielle Dionne, Sarah Hopp, Matthew P. Frosch, Bradley T. Hyman, Thomas Beach, Rani Sarkis, Garth R Cosgrove, Jeffrey Helgager, Jeffrey A. Golden, Page B. Pennell, Julie A. Schneider, David A. Bennett, Aviv Regev, Wassim Elyaman, Elizabeth M. Bradshaw, Philip L. De Jager

## Abstract

Recent studies of bulk microglia have provided insights into the role of this immune cell type in central nervous system development, homeostasis and dysfunction. Nonetheless, our understanding of the diversity of human microglial cell states remains limited; microglia are highly plastic and have multiple different roles, making the extent of phenotypic heterogeneity a central question, especially in light of the development of therapies targeting this cell type. Here, we investigated the population structure of human microglia by single-cell RNA-sequencing. Using surgical- and autopsy-derived cortical brain samples, we identified 14 human microglial subpopulations and noted substantial intra- and inter-individual heterogeneity. These putative subpopulations display divergent associations with Alzheimer’s disease, multiple sclerosis, and other diseases. Several states show enrichment for genes found in disease-associated mouse microglial states, suggesting additional diversity among human microglia. Overall, human microglia appear to exist in different functional states with varying levels of involvement in different brain pathologies.

## Introduction

Our understanding of microglia has evolved rapidly with respect to their ontology, role in developmental and physiological plasticity, and involvement in pathophysiology^1^,^2^. Historically, morphological studies using tissue sections^3^ led to the notion of a linear continuum of cell states from a homeostatic state with a ramified morphology to an activated state with globular morphology. Recent transcriptome-wide studies of bulk *ex vivo* human microglia have consistently suggested that microglia change with age and have profile senriched for disease genes^4^^-^^6^. Further, we observed that one microglial transcriptional program contributes to the accumulation of tau pathology while two others may relate to β-amyloid pathology^7^. However, these analyses used cortical-level data, and the need for greater resolution led us to characterize heterogeneity of human microglia at the single-cell level.

Genetic studies have highlighted a prominent role for microglia in susceptibility to different neurodegenerative diseases, particularly Alzheimer’s disease (AD) and multiple sclerosis (MS)^6^,^8^. These syndromic diseases are heterogeneous at the clinical and pathologic level; for example, in a selection of over 1000 older individuals, more than 230 unique combinations of neuropathologic features were observed^9^. These concurrent pathologic processes create a diversity of contexts to which microglia respond, suggesting there may be large inter-individual variation in microglial states layered onto topological and functional variation of homeostatic microglia. This diversity of states makes targeting microglia in disease challenging: one has to carefully map which microglial subset to modulate and in which direction the immunomodulation must be applied.

To address this diversity, we profiled 15,910 cells isolated from the cerebral cortices of 7 aged and 8 young and middle-aged adults. Within the microglial cells (>97% of the cells), our data identified substantial heterogeneity in cell states, yielding an initial catalog of human microglial subpopulations. Compared to previous reports on mouse microglia^10^,^11^, the human brain seems to harbor a more nuanced impact of different disease processes that suggest an element of specialization of microglial cell states to different pathologic contexts.

## Results

### Nature and distribution of human single cell transcriptomic data

We elected to include a variety of adult human subjects to ensure that we captured a reasonable diversity of cortical microglia. **Supplementary data 1** outlines the demographic characteristics and provenance of each of the profiled samples. In short, we processed 7 fresh autopsy samples from the dorsolateral prefrontal cortex, as well as 2 fresh samples of hippocampus and 6 samples of temporal neocortex from epilepsy surgeries. All samples were processed in the same manner, as previously described^6^, to extract live microglia: we sorted all CD45+/CD11b+/7AAD- cells from the autopsy samples (**Figure 1a**). In addition, since we observed the presence of rare CD45+/CD11b- events in the surgical biopsy samples, these were also included in the sequencing effort to comprehensively capture all viable immune cells in the CNS. The purified cell suspension from each processed sample was profiled using the droplet-based Chromium platform from 10x Genomics (Methods).

**Figure 1.**
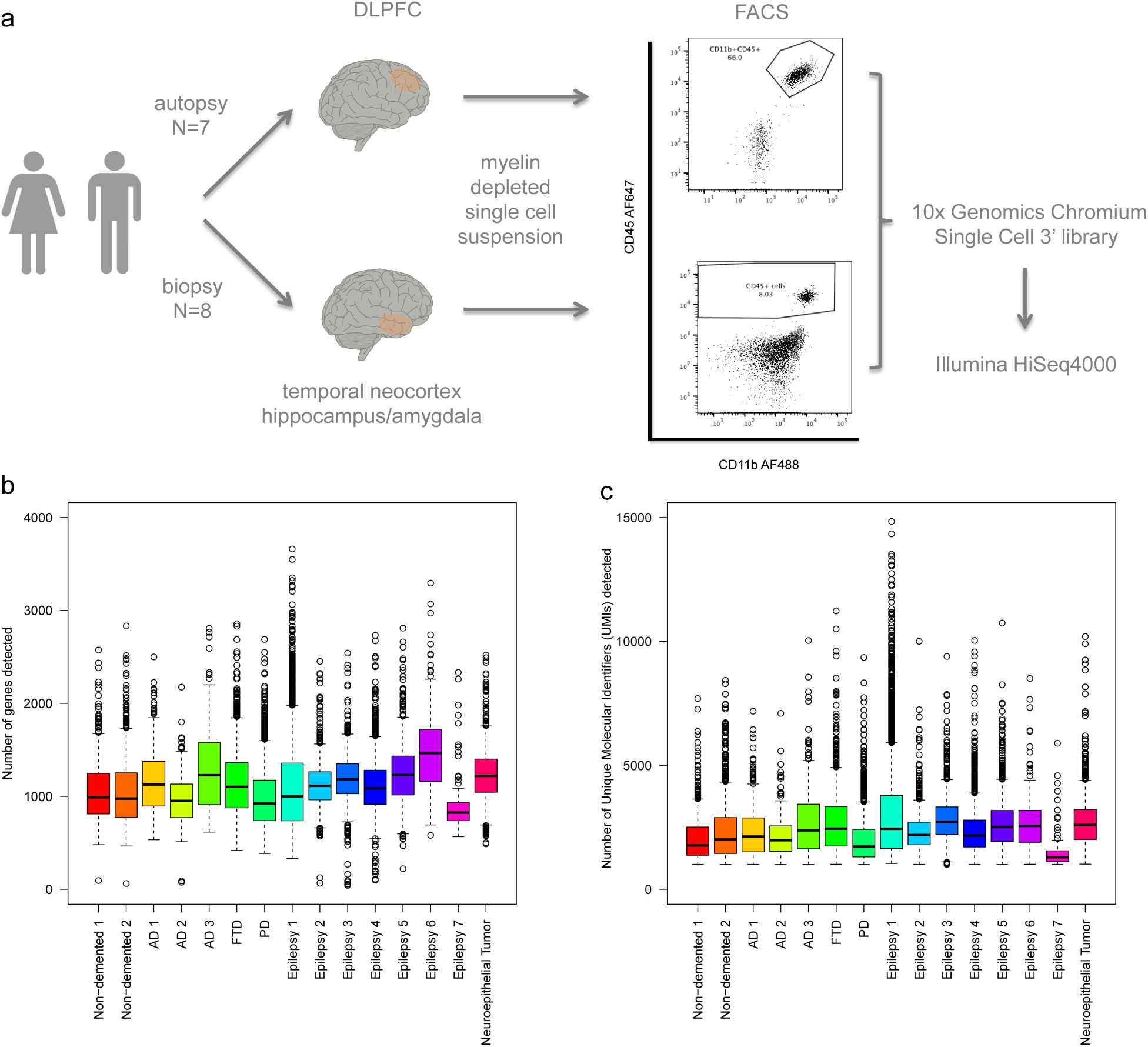
Workflow of human brain myeloid cell scRNAseq study and quality control of the resulting dataset. **(a)** Workflow. Brain myeloid cells were isolated from 15 donors of both sexes (for a detailed isolation protocol see ^6^). Autopsy samples originated from the DLPFC of deceased aged individuals with various pathologies, while surgical biopsy samples were from the temporal neocortex and hippocampus of young and middle-aged individuals undergoing temporal lobectomy surgery for intractable epilepsy. Brain myeloid cells were sorted based on their CD11b and CD45 expression in autopsy samples. In surgical samples, a CD45 single positive population was observed and was included in the purified sample. Each single cell suspension preparation of sorted brain myeloid cells was loaded onto one lane of the Chromium system (10x Genomics) and the resulting library was sequenced on the HiSeq4000 platform (Illumina). **b-c** Quality control of the scRNAseq dataset. The box plots represent the median (middle line), 25%, and 75% percentiles. Whiskers extend to the most extreme cell no more than 1.5 times the interquartile range, and all cells more extreme than the whiskers are represented as circles. Abbreviations: AD, Alzheimer’s disease; AF647 (AlexaFluor647) and AF488 (AlexaFluor488), two different fluorochromes; DLPFC, dorsolateral prefrontal cortex; FACS, fluorescence activated cell sorting; FTD, fronto-temporal dementia; PD, Parkinson’s disease; scRNAseq, single cell RNA sequencing; UMI, unique molecular identifier.

A rigorous pre-processing pipeline (Methods) yielded 15,910 individual cells with a median of 1,246 cells sequenced per subject (**Supplementary data 2**). The mean number of Unique Molecular Identifiers (UMIs) and genes detected per cell in each subject (**Figure 1b,c**) was comparable among the different donors and specimen (autopsy vs surgical) types. We ran an iterative PCA-Louvain clustering^12^,^13^ approach with stepwise cluster robustness assessment and identified 23 distinct cell clusters with a minimum of 20 cells per cluster (**Figure 2a**). The number of UMIs and the number of detected genes was comparable among the different clusters (**Supplementary figure 1**).

**Figure 2.**
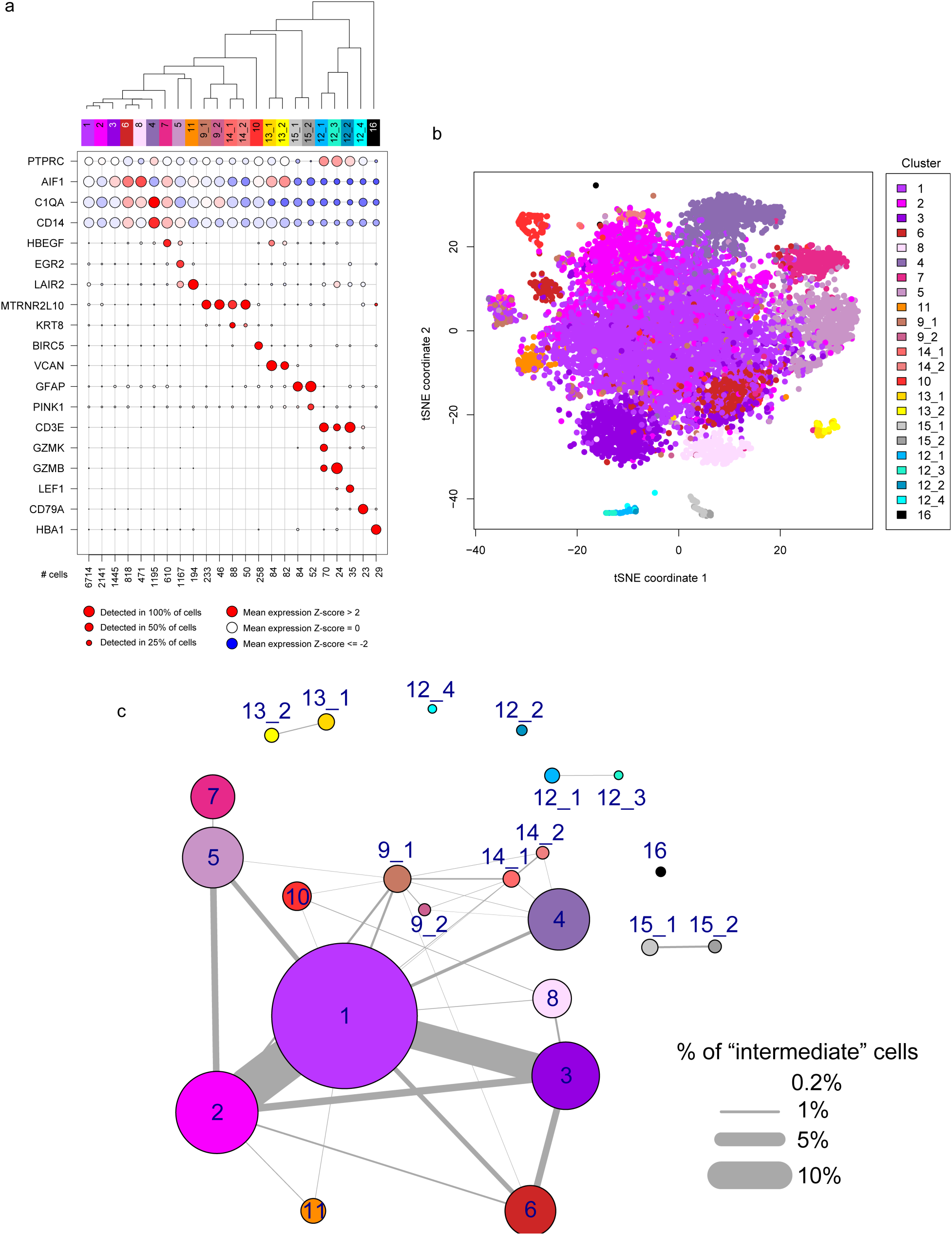
scRNAseq identifies unique subsets of human brain myeloid cells. **(a)** Unsupervised iterative PCA-Louvain clustering with a random forest cluster robustness assessment identified 23 different clusters of cells in our dataset. Each column represents a cell cluster; they are ordered in relation to similarities in their gene expression patterns. The number of cells assigned to each cluster is noted at the bottom of each column. In rows, we represent the level of expression of selected key genes. The size of the dot represents the fraction of cells in a given cluster in which the gene was detected (>0 counts per million). The color of the dot represents the average expression z-score (calculated over all 15,910 cells) of the cells within a given cluster. The bulk of the cells belonged to 16 clusters identifiable as myeloid based on their marker gene expression (AIF1/IBA1 and CD14). Of these, 2 clusters (13_1 and 13_2) had low expression of C1QA, a microglia marker and probably represent monocytes. A small proportion (~2%) of the cells were non-myeloid and belonged to clusters that could be characterized by high expression of genes such as GFAP (cluster 15_1 and 15_2), CD3E (cluster 12_1, 12_2, 12_3), CD79A (cluster 12_4) and HBA1 (cluster 16), likely representing astrocytes, T cells, B cells and erythrocytes, respectively. The dendrogram at the top represents the hierarchical clustering using the mean expression profile of each group. **(b)** t-SNE plot of the assayed cells. Each dot represents a cell, which is color coded based on its cluster identity. The color key is displayed to the right of the panel. There is clear segregation of the non-microglial clusters (the monocytic cluster in yellow (13_1 and 13_2), the astrocytic cluster in grey (15_1,15_2), the T/B cell cluster in teal (cluster group 12), and the erythrocyte cluster in black (16) from the microglia clusters, to which the bulk of the cells belong. **(c)** Constellation diagram showing the relationship between the different clusters. For every pair of clusters, a bootstrapped random forest approach was run to classify each cell 10 times. Each node represents a cluster, scaled by the number of cells that belong to it, and each edge represents the fraction of cells that are not 100% classified into a single cluster for each pair of clusters. The largest cluster (cluster 1) shares substantial “ambiguously classified cells” with clusters 2 and 3, which may suggest a continuum of states among these three clusters. The other microglial clusters share fewer cells with cluster 1, with varying level of connectivity, while the monocyte and non-myeloid clusters have no edge with the microglial clusters. Abbreviations: tSNE, t-Distributed Stochastic Neighbor Embedding.

In 16 out of 23 clusters (>98% of cells), we detect known myeloid markers (CD14, AIF1/IBA1) (**Figure 2a**). The remaining cells are distributed among a set of putative T-cell clusters (12_1 through 12_3), a B cell cluster (12_4), and two minor ambiguous clusters (15_1 and 15_2) expressing myeloid markers (such as AIF1/IBA1) as well as high levels of GFAP (**Figure 2a**), MBP and SNAP25 (data not shown). These latter two clusters could comprise cell doublets, but we cannot unambiguously call them as such based on numbers of genes or UMIs detected (**Supplementary figure 1**). Additionally, we detect 29 cells (cluster 16) that are likely erythrocytes, based on hemoglobin expression. Among the 16 myeloid clusters, we found that clusters 13_1 and 13_2 expressed (1) C1QA, a microglial marker, at very low levels and (2) high levels of monocyte specific genes, such as *FCN1*, *VCAN*, and *LYZ* (**Supplementary figure 2a**), suggesting these two clusters may represent monocytes or monocyte-derived cells. By contrast, the remaining 14 clusters express high levels of microglia-specific genes, such as *C1QA*, *C1QB*, *C1QC* and *GPR34* (**Supplementary figure 2b**); we therefore deem these 14 clusters to be distinct clusters of microglial cell states.

Visualization of all the profiled cells in a t-SNE projection (**Figure 2b**) shows that the non-microglial clusters (13_1, 13_2, 15_1, 15_2, 12_1, 12_2, 12_3, 12_4 and 16) segregate from the microglial clusters. We assessed inter-cluster relatedness using a random forest-based machine learning approach to characterize how well individual cells could be unambiguously classified to each cluster^14^ (see Methods section). We generated a “constellation diagram”^15^ representing the likelihood of each cell belonging to any cluster in a given pair (**Figure 2c**). The top three clusters - accounting for 65% of cells – have a large proportion of cells ambiguously classified among them. This interrelatedness among clusters 1, 2, and 3 suggests that they comprise cells in closely related transcriptomic states. The remaining microglial clusters show more distinct signatures, as evidenced by the smaller proportion of ambiguous cells. Interestingly, this analysis does not support the concept of a single linear relationship among clusters; rather, independent microglial clusters appear to share similarities with cluster 1 but less so with each other. Although RNA-seq data alone is not sufficient to distinguish cell subtypes from cell states, the degree of similarity between clusters 1, 2, and 3 suggests that they might be different states of the same underlying cell subtype, whereas the remaining clusters might represent more specialized subtypes of microglia. Finally, there are no ambiguous cells between any of the non-microglial and microglial clusters, highlighting the clear difference between microglia and other cell classes.

We then assessed the extent of inter-individual heterogeneity in our data (**Supplementary figure 3**). First, cluster 1 and 2 are well-represented in most individuals (**Figure 3a**). We therefore propose that cluster 1 may represent “homeostatic microglia” which fulfill routine tasks necessary for cortical function. Second, there is inter-individual variability in the frequency of other microglial clusters, and a subset of clusters (9_1,9_2,14_1,14_2) are present in only two older individuals (95 and 97 years old) or a single subject with epilepsy (cluster 11) (**Figure 3b**). Further, one subject, “Epilepsy 1”, has a different cluster frequency pattern from all other subjects (**Figure 3a**): clusters 1 and 2 are relatively small and the majority of microglia are distributed across 7 other clusters. For this subject alone, the tissue sample was taken from cortex that was monitored using subdural electrodes placed for intracranial electroencephalogram monitoring for nine days prior to the resection. The resulting distribution of microglia away from cluster 1 could thus represent a relatively nonspecific reaction away from homeostasis, caused by the presence of a foreign object. Interestingly, clusters 9_1, 9_2, 14_1 and 14_2 were not induced in this sample, suggesting that they may not represent a non-specific reactive phenotype but might rather be an age-related state.

**Figure 3.**
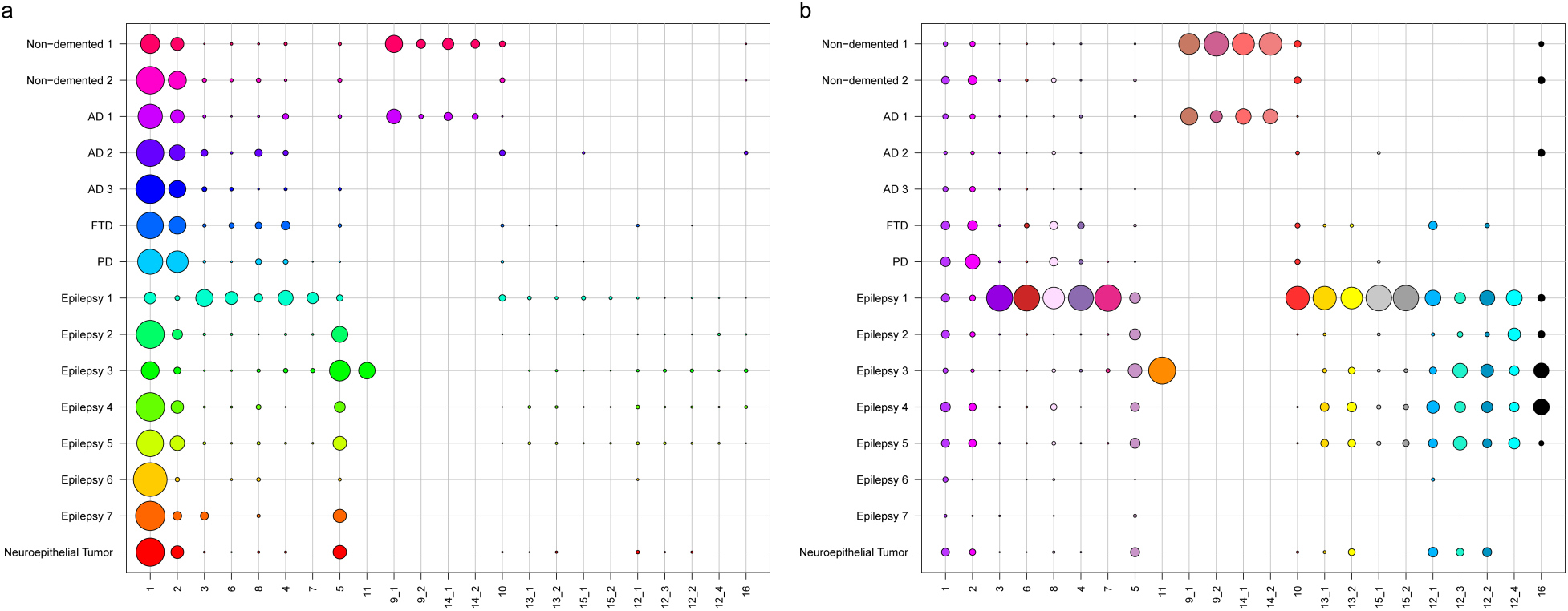
Cluster distribution within donors and provenance of clusters. **(a)** Distribution of the different cell clusters within donors. Each column represents a cluster, and each row represents a subject. The size of the circles corresponds to the number of cells present in each cluster for each subject. Cluster 1 is the largest cluster in most subjects. The data is normalized by rows. In each row, the different circles add up to 100%. Each subject is colored with a different color. **(b)** Representation of the subject-level origin of the cells in each cluster. The size of the circles corresponds to the number of cells assigned to each cluster in that subject. The data is normalized by columns so that the different circles add up to 100% in each column. Thus, for example, while most subjects have cells assigned to Cluster 6, the vast majority of cells assigned to Cluster 6 came from subject “Epilepsy 1.” Circles are colored by clusters, following the color code outlined in **Figure 2**.

### Annotating the clusters of human microglia

We next examined genes showing expression restricted to specific microglial subtypes. Unique sets of transcription factors and transcriptional regulators were found in clusters 5, 7, 10, 11, 9_1, 9_2, 14_1, and 14_2, but not in clusters 1, 2, 3, 4, 6, and 8 (**Figure 4a**). The same is true for cell surface markers (**Figure 4b**), with the exception of cluster 4. The lack of distinct on-off transcription factors and cell-surface markers among the first three clusters (1, 2, and 3) is consistent with our hypothesis that these three clusters represent homeostatic microglia from which the other clusters differ by the up-regulation of specific genes. From a global perspective (**Figure 4c and d**), cluster set 14 has the largest number of differentially expressed transcription factors and cell surface molecule-encoding genes, whereas cluster 10 showed upregulation of transcription factors but not cell surface molecule-encoding genes. For a subset of clusters with distinct markers (**Figure 5a)**, we confirm that the corresponding protein expression is restricted to a subset of microglia in the human brain using immunofluorescence staining on tissue sections (**Supplementary figure 4**). The other clusters lack such a unique signature but can be defined by combinations of on-off genes or differing expression levels of multiple genes. For example, clusters 1, 2, and 3 are distinguished from one another by a gradient of expression among multiple genes (**Figure 5b**).

**Figure 4.**
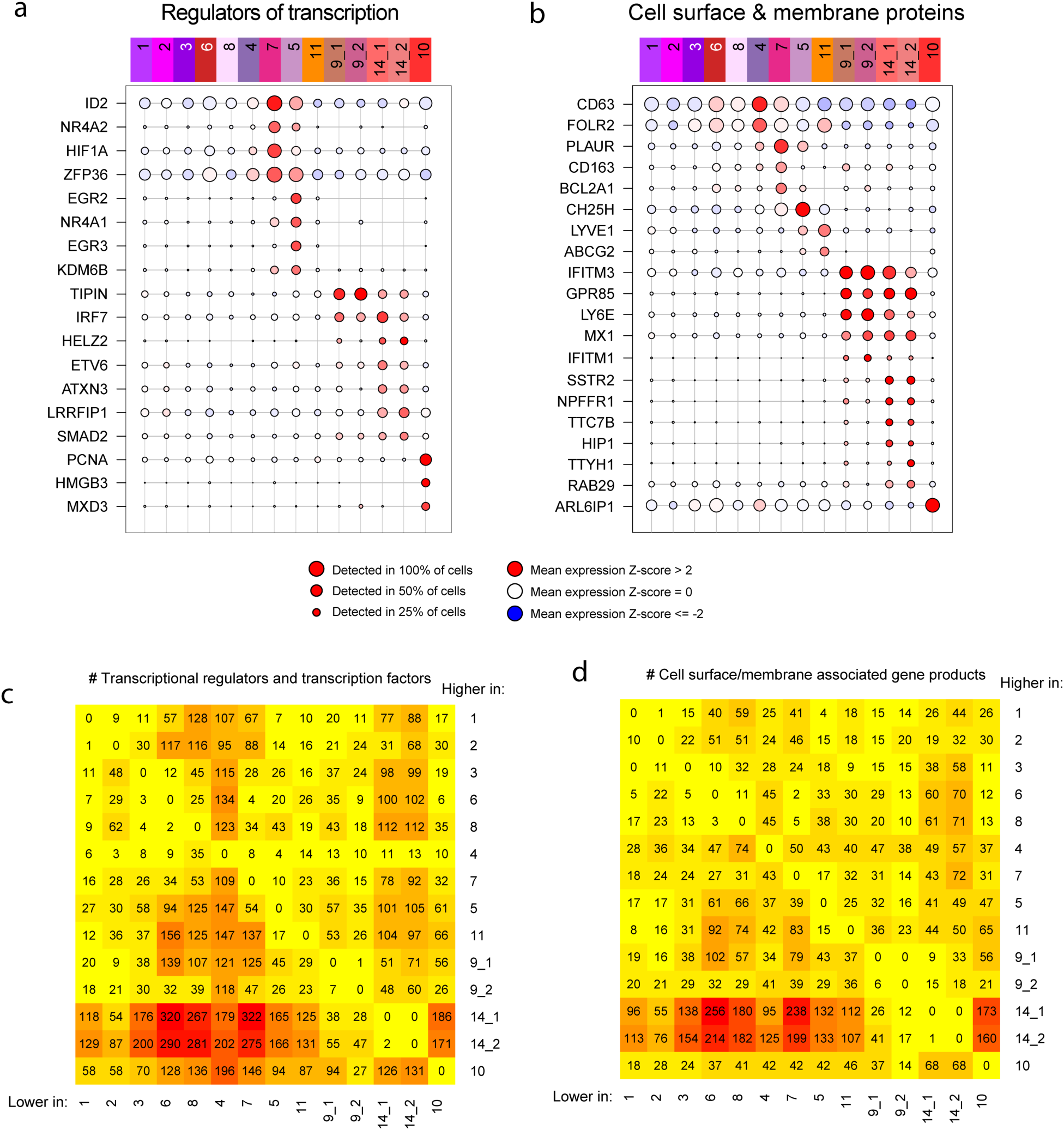
Identifying potential marker genes for the microglial clusters. **(a)** Microglial clusters are visualized in columns, and selected transcription factors that are enriched in certain clusters are listed in rows. The key code at the bottom of the panel illustrates the fact that the size of each dot represents the fraction of cells in a given cluster in which the gene was detected (>0 counts per million), and the color of the dot represents the mean of the expression z-score (calculated using all 15,910 cells) for the cells belonging to that cluster, as in Figure 2. **(b)** Same as a), but for cell surface genes that are differentially expressed across clusters, which make good targets for antibody development. (**c-d)** Heatmaps representing the number of differentially expressed genes in each pairwise comparison between the microglial clusters. In **(c)**, we limit the analysis to genes that encode transcription factors and transcriptional regulators. In **(e)**, we present the results of an analysis limited to genes encoding cell surface proteins.

**Figure 5.**
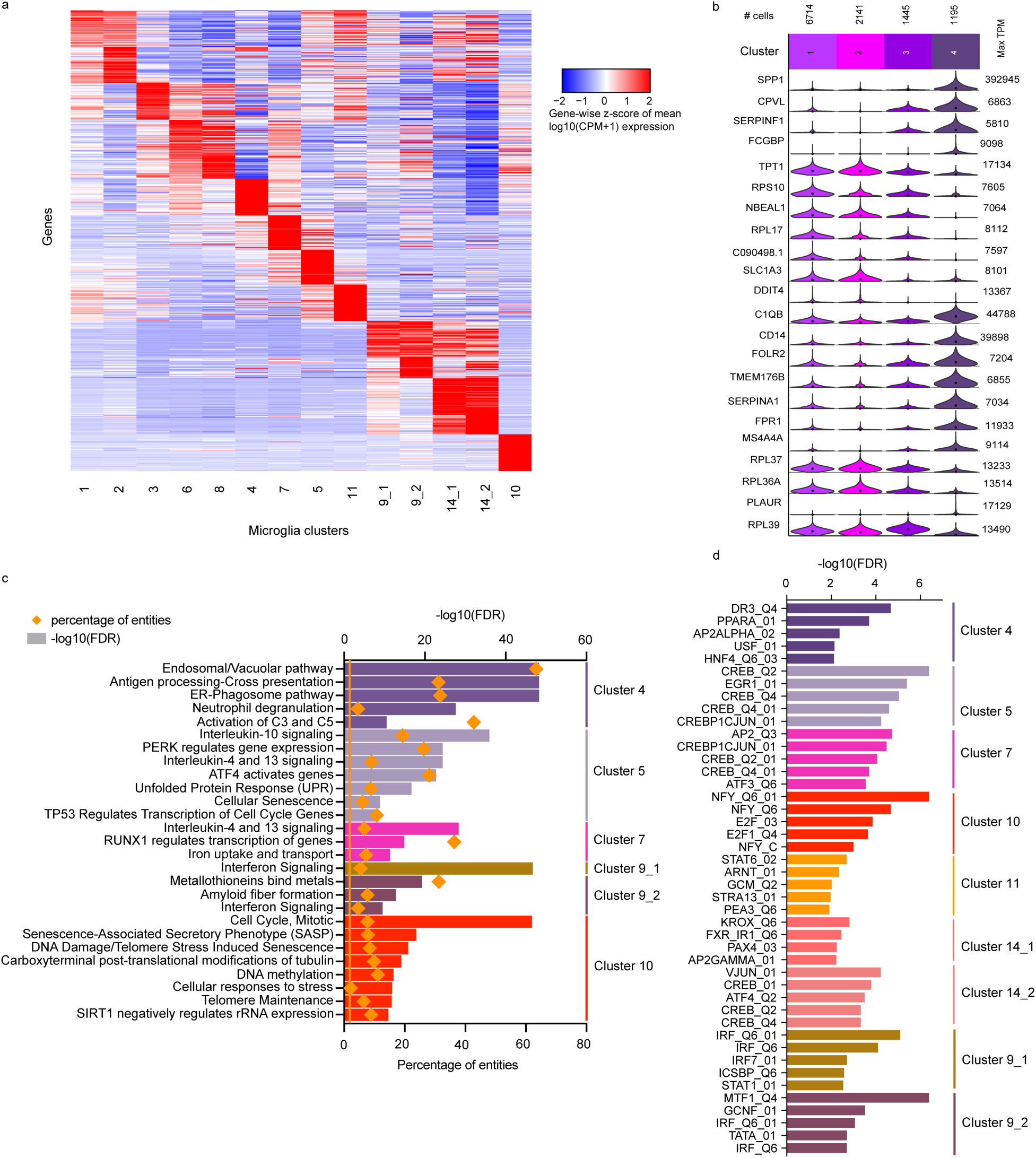
Functional annotation of the microglial clusters. **(a)** Heatmap depicting the top 20 signature genes of each microglial cluster. Rows represent genes, and each cluster is presented in a column. The color coding represents the mean expression over the cluster, Z-scored over all the clusters. **(b)** Violin plot showing the gene expression differences between the first four clusters (in columns) for selected genes (in rows) that illustrate the relative differences between these four clusters. These four microglia clusters contain the majority of cells and are governed by gradients in the expression levels of genes such as *SPP1* and *MS4A4A*. **(c)** Functional annotation of certain microglia clusters using REACTOME pathways enriched for their signature genes (top 50 differentially expressed genes). Pathways are color coded by cluster, and clusters are listed to the right of the panel. **(d)** Predicted transcription factors whose binding sites are enriched among the component genes of each microglial cluster. In rows, the names of the consensus transcription binding site matrices are shown. Enrichment p values were calculated with PASTAA. The transcription factors are color coded by cluster, and the clusters are listed to the right of the panel. Abbreviations: TPM, transcripts per million; FDR, false discovery rate.

For functional annotation, we focused on those clusters for which a unique set of enriched genes could be identified. Gene set analysis^16^ (**Figure 5c, Supplementary data 3**) reveals that cluster 4 is enriched in genes related to antigen presentation, while cluster 5 and 7 feature genes related to anti-inflammatory responses (IL-10, IL-4 and IL-13). Cluster 9 is enriched in genes belonging to the interferon signaling pathway, and cluster 10 is enriched in genes associated with the cell cycle, suggesting that it may constitute a pool of proliferating microglial cells. We extended this analysis by assessing which transcription factor binding sites are enriched in the promoters of differentially expressed genes among these clusters. Using the PASTAA software^17^, some of the strongest enrichments are observed for *CREB* in cluster 5, *RUNX1* in cluster 10 and IRF transcription factors and *STAT1* regulated genes in cluster sets 9 and 14 (**Supplementary data 4**), prioritizing regulators that may play an important in each microglial subset.

We then turned to the annotation of our microglial clusters using other signatures found in the literature. In a recent study of the CKp25 mouse model, a microglial subset enriched in interferon response genes was implicated in late microglial responses to this *in vivo* perturbation, and a different subset was implicated in the early response^10^. In our data (**Supplementary figure 5**), the mouse early response genes were only detected in cluster 10, suggesting that the early microglial response in the CKp25 mouse model may involve a proliferative reaction. The late response signature appears more nuanced in humans, with component genes from the mouse study being found in either all human microglial clusters or limited to cluster sets 9 and 14, in line with the enrichment of these two sets of clusters in interferon response pathways (**Supplementary data 4**). As mentioned above, these four clusters are found only in 2 aged individuals, suggesting that they have a very context-specific function. Similarly, a meta-analysis of all of the currently available mouse microglia RNAseq datasets determined gene co-expression modules with unique functionality^18^. Here again, we find a co-expression module that captures the putative proliferating microglia of cluster 10 (**Supplementary figure 6a)** and an interferon response module (**Supplementary figure 6b)**. The modules relating to LPS response (**Supplementary figure 6c)** or neurodegeneration in this analysis of murine data (**Supplementary figure 6d)** were not enriched in our human clusters.

We also specifically evaluated our human clusters in regards to a recent report of a “disease-associated microglia” (DAM) signature^11^ (**Figure 6** and **Supplementary figure 7).** Most of the DAM genes were detected in multiple microglial clusters (**Figure 6a, 6b** and **Supplementary figure 7a and 7b**). Correlations among DAM genes were weak throughout the data set, with the exception of *P2RY12* and *APOE,* which display the expected anti-correlation in expression levels. We found that cluster 4 showed the strongest enrichment for the DAM expression profile, with clusters 6, 8, 9_1, and 9_2 also showed higher enrichment for the DAM expression profile than the non-DAM expression profile (**Figure 6b, Supplementary data 5**). Thus, while cluster 4 may be the most DAM-like cluster, the function attributed to DAM in mice is more distributed in human microglia. This observation probably has multiple contributing factors, including the phenotypic diversity of human subjects, the potential specialization of different microglial subsets to distinct contexts in humans, and technical factors that may have reduced the resolution of the murine data.

**Figure 6.**
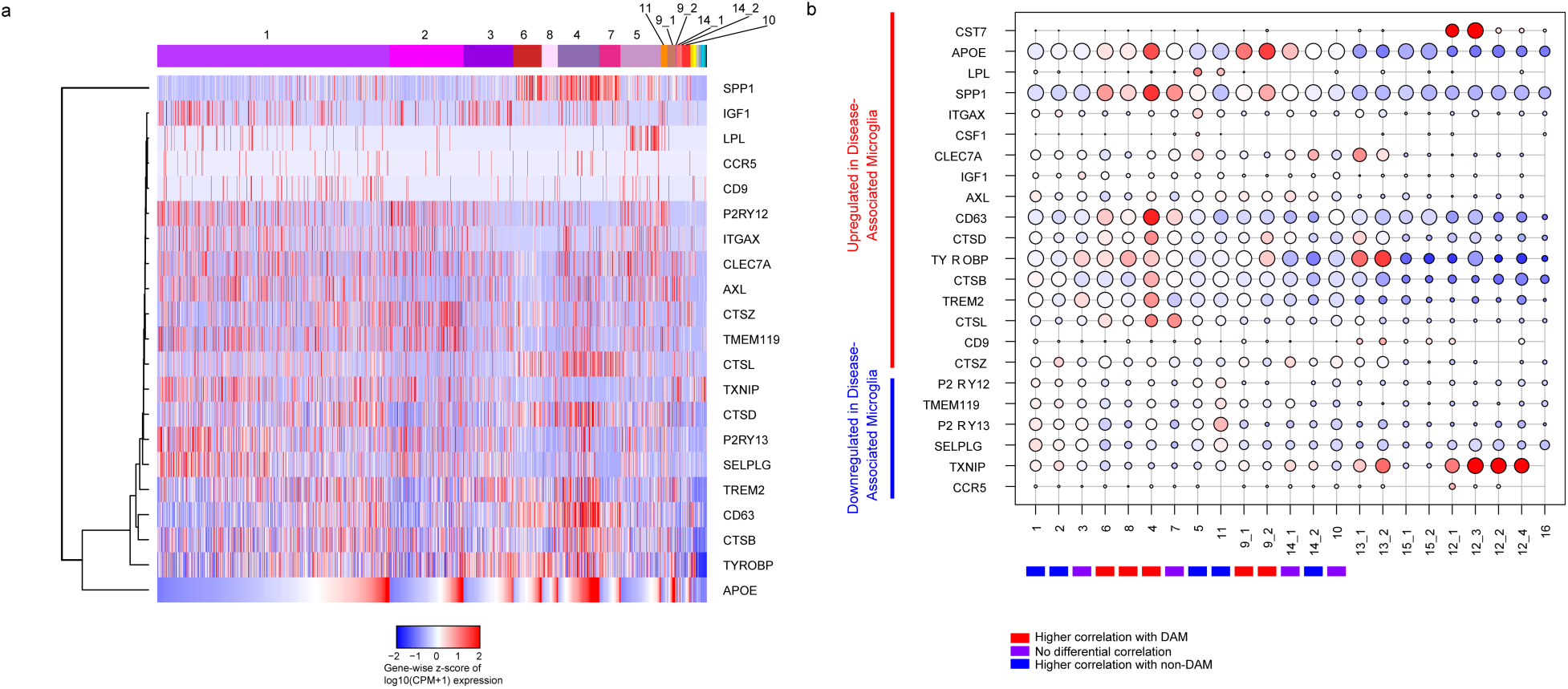
Investigation of the murine DAM phenotype in human microglia. **(a)** Heatmap depicting the expression levels of the genes in the murine Disease Associated Microglia (DAM) gene set. Each column represents a cell. Cells are ordered based first on cluster and then APOE expression within each cluster. The clusters are labeled at the top of the panel. Genes (rows) are ordered based on unsupervised hierarchical clustering. The color code represents Z-score of expression for each gene (i.e. normalized by row). While some of the DAM genes show some correlation in expression levels across cells, the gene set does not appear to be as coherent as it is in mice. **(b)** Figure depicting the expression levels of genes related to the DAM phenotype in each microglial cluster. Each cluster is found in one column. The selected genes are either upregulated (upper set of genes) or downregulated (lower set of genes) in DAM. The size of the dots is proportional to the number of cells expressing the given gene in the corresponding cluster. The color of the dots represents the mean Z score of expression. Bars on the bottom indicate whether the mean cluster expression correlated more strongly with the DAM (red) or the homeostatic (blue) murine microglia RNA-seq expression profile (from Keren-Shaul et al. 2018). Clusters 6, 8, 4, 9_1 and 9_2 showed the strongest enrichment for the DAM phenotype, while clusters 1, 2, 5 and 11 were enriched for the homeostatic signature which is downregulated in DAMs.

### Discrete microglial subsets are enriched for MS and AD genes

Given that we have too few subjects to directly evaluate the association of microglial clusters to diseases and human traits, we performed enrichment analyses (Methods) to identify clusters that may be implicated in disease. We used the DOSE Bioconductor package^19^ to assess statistical enrichment for disease-related genes (**Fig 7a, Supplementary data 6**). The most striking enrichment is seen for cluster 4 and amyloid and Lewy body pathology as well as multiple sclerosis. Cluster 4 also has a more modest enrichment for tauopathy-related genes, as do the peripheral myeloid clusters (13_1 and 13_2, **Supplementary figure 8**), while cluster 5 shows some enrichment for motor neuron disease genes. We also see that many individual GWAS identified risk genes for AD, MS, PD and ALS (**Supplementary figure 9a** through **d**, respectively) are expressed in most of the clusters, with a subset strongly expressed in cluster sets 9 and 14. Interestingly, the TSPO gene, the target for all current microglial markers used in positron emission tomography studies, is expressed in all clusters (**Supplementary figure 9e**) and is therefore a good proxy for total microglial count.

**Figure 7.**
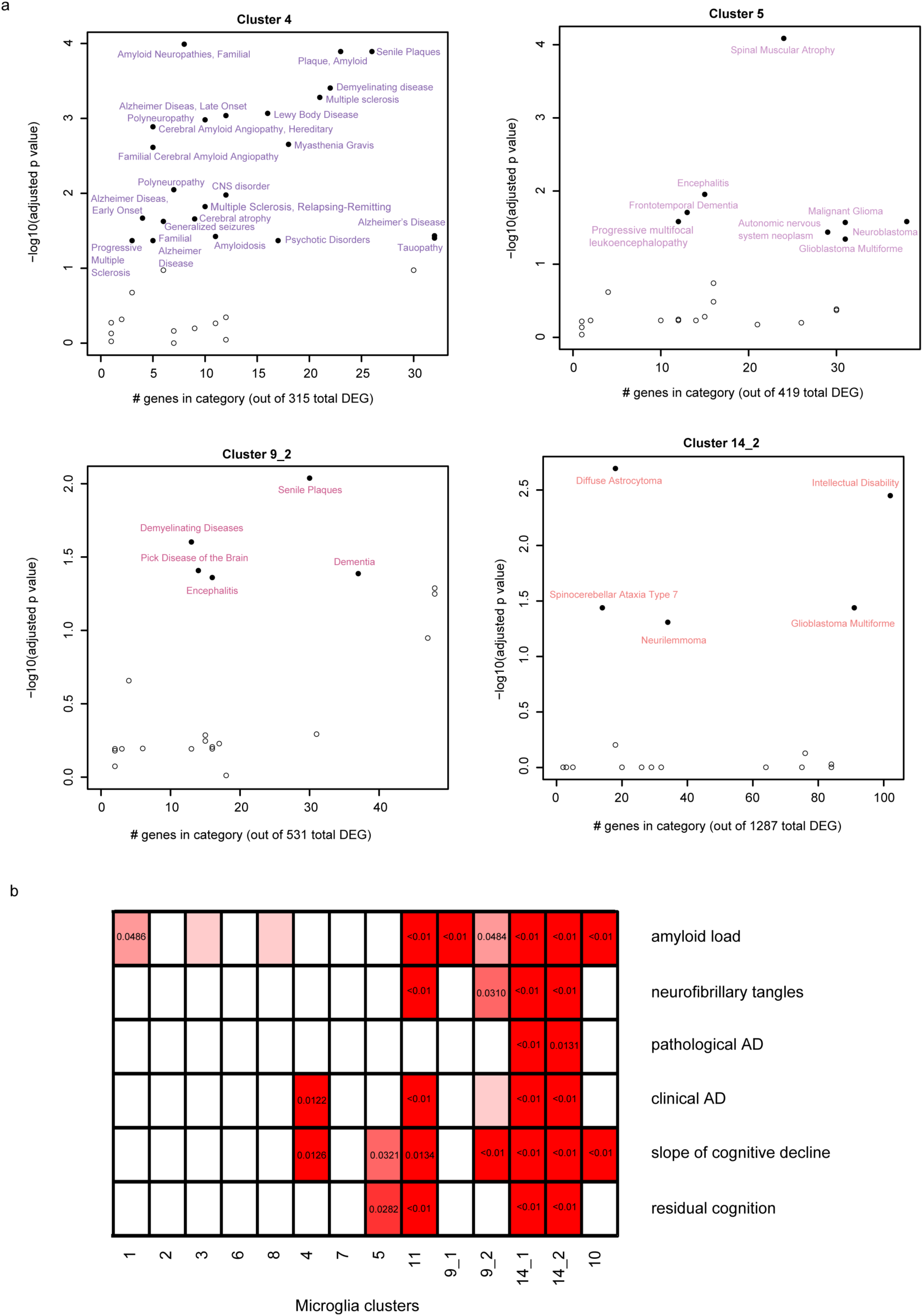
Disease association in human microglia clusters. **(a)** Scatter plots depicting brain related diseases – using gene sets from the disease ontology database (http://disease-ontology.org/) - that are significantly enriched (adjusted p-value <0.01, hypergeometric test with BH correction) in a given microglia cluster - using the signature gene sets of each microglia cluster. Results for four different clusters are shown; the other clusters are included in **Supplementary figure 7**. In each plot, the y-axis reports the p-value of the enrichment analysis while the x-axis reports the number of genes which overlap between the cluster and disease gene sets. **(b)** Panel reporting the result of enrichment analyses between the genes defining the microglia clusters and those genes that are associated with pathological or clinical traits found in the aging human brain. Significant p-values are shown for those cluster/trait combination where they are significant, and the color of each box is related to the strength of the association. The adjusted p-value shown here are obtained from the rank-rank hypergeometric test. Abbreviations: AD Alzheimer’s disease.

We also used a rank-rank hypergeometric approach^20^ to explore the relationship between the different microglia clusters and aging-related traits (**Figure 7b, Supplementary data 7**). Since many of our samples came from autopsies of older individuals, we assessed for enrichment of gene signatures derived from our analyses of cortical tissue RNAseq profiles in 541 individuals in each set of cluster-specific upregulated genes^21^. We evaluated genes associated with either a clinical or pathologic diagnosis of AD as well as with (1) the molecularly specific measures of β-amyloid and PHF-tau tangle accumulation, (2) the slope of cognitive decline before death and (3) a measure of resilience to cognitive decline (“residual cognition”)^22^,^23^. We find that clusters 14_1 and 14_2 (which contain genes related to the interferon response) are enriched for all of the investigated traits, while cluster 4 only showed an enrichment for Alzheimer’s dementia and cognitive decline but not with the histopathological hallmarks of the disease (**Figure 7b**).

We also evaluated modules of co-expressed genes defined in the aging human frontal cortex^21^ that we had previously described as being enriched in microglial genes^7^. These five modules include m116, the cortical module most enriched for microglial genes, and m5, which is associated with both accumulation of tau pathology and the number of morphologically activated microglia in cortical tissue^24^. Gene-set enrichment analysis (**Supplementary figure 10, Supplementary data 8)** shows that m116 is enriched in almost all microglial clusters while m5 is present in a subset of cell clusters. This result suggests that morphologically “activated” microglia may exist in different transcriptomic states of activation. That each of these five microglial modules defined in tissue-level cortical RNAseq data are enriched in a majority of clusters is not surprising, as only those signatures shared by a large number of microglia will emerge in tissue level data. The important corollary to this point is that tissue-level data, while rich in many respects, is inadequate for the detailed investigation of the role of microglia in neurodegenerative diseases and aging.

## Discussion

This manuscript presents a new **Resource** based on 15,430 individual human microglial transcriptomes (from 15,910 total number of profiled cells) which are derived from 15 individuals and identify 14 putative clusters of microglia. Our analysis identifies the microglia subsets involved in homeostasis, proliferation, interferon response, and diseases such as AD and multiple sclerosis. We document inter- and intra-individual heterogeneity in microglial states, including intriguing subpopulations with an interferon response present only in 2 older individuals. Our data are searchable at https://vmenon.shinyapps.io/microglia.

This report contains several insights. First, the major populations of microglia are found in both autopsy and surgical samples, and their frequencies are similar across both types of samples. This suggests that these two sources of primary, live microglia do not have large differences arising from technical factors or circumstances surrounding the agonal state. We noted the presence of peripheral lymphoid cells in the surgical samples and therefore gated on all CD45+ cells for these samples. The total number of non-microglial cells is very small, and their provenance is unclear (blood *vs.* infiltrating cells). Second, multiple clusters contain disease-related genes. Thus, the role of microglia in human disease is likely to be more nuanced than what has been described in the mouse to date, but cluster 4 stands out amongst the other clusters as it emerges as disease-related from multiple different analyses. Finally, we find that several clusters which are enriched in interferon response genes are only seen in two older subjects and another (cluster 11) is seen only in one epilepsy subject, suggesting that there may be additional microglial states to discover in other human samples.

Our study has certain limitations that result from having profiled samples from a small number of individuals, given the difficulty in obtaining these samples and the use of a multi-step purification pipeline^6^ to isolate live microglia from human brain tissue. First, there may have been a survival bias among microglial subpopulations that are successfully profiled. Further, our sample preparation process results in the loss of potentially important topological information. Also, we have only sampled three cortical regions, and thus profiling a larger number of brain regions and subjects is necessary to improve our reference atlas. Finally, all of the disease analyses reported here are indirect, relying on enrichment analyses, and they will need to be confirmed by direct analysis as larger sample sizes become available.

Overall, this **Resource** opens several avenues of investigation: (1) the exploration of functions conducted by the human homeostatic microglial subtype that we have now defined molecularly, (2) the generation of more complete transcriptomes, epigenomes, and proteomes to elucidate the function of each cluster now that we have markers with which to purify them, (3) enhanced *in silico* analyses of genetic, or tissue-level transcriptomic and epigenomic data to assess which microglial subtypes are involved in traits of interest, and (4) the identification of the specific subset of microglia to prioritize in target validation and clinical development efforts that will lead to therapies that modulate microglial function in neurodegenerative diseases.

## Data availability

The single cell based transcriptomic atlas of human microglia presented here is available in the form of a searchable platform at https://vmenon.shinyapps.io/microglia. The raw data files are available through Synapse (https://www.synapse.org/#!Synapse:syn3219045).

## Acknowledgements

We thank the individuals who have generously donated samples of their brain to research either through the RUSH University Alzheimer’s Disease Center, the Brigham & Women’s Hospital, the Massachusetts Alzheimer’s Disease Research Center, or the Banner Sun Health Research Institute. This research is supported by a fellowship from Alzheimer’s Association and by grants from the National Institute of Aging (U01 AG046152, R01 AG048015, RF1 AG057473) and the National Institute for Neurologic Disease and Stroke (R01 NS089674).

## Figure legends

**Supplementary figure 1.**
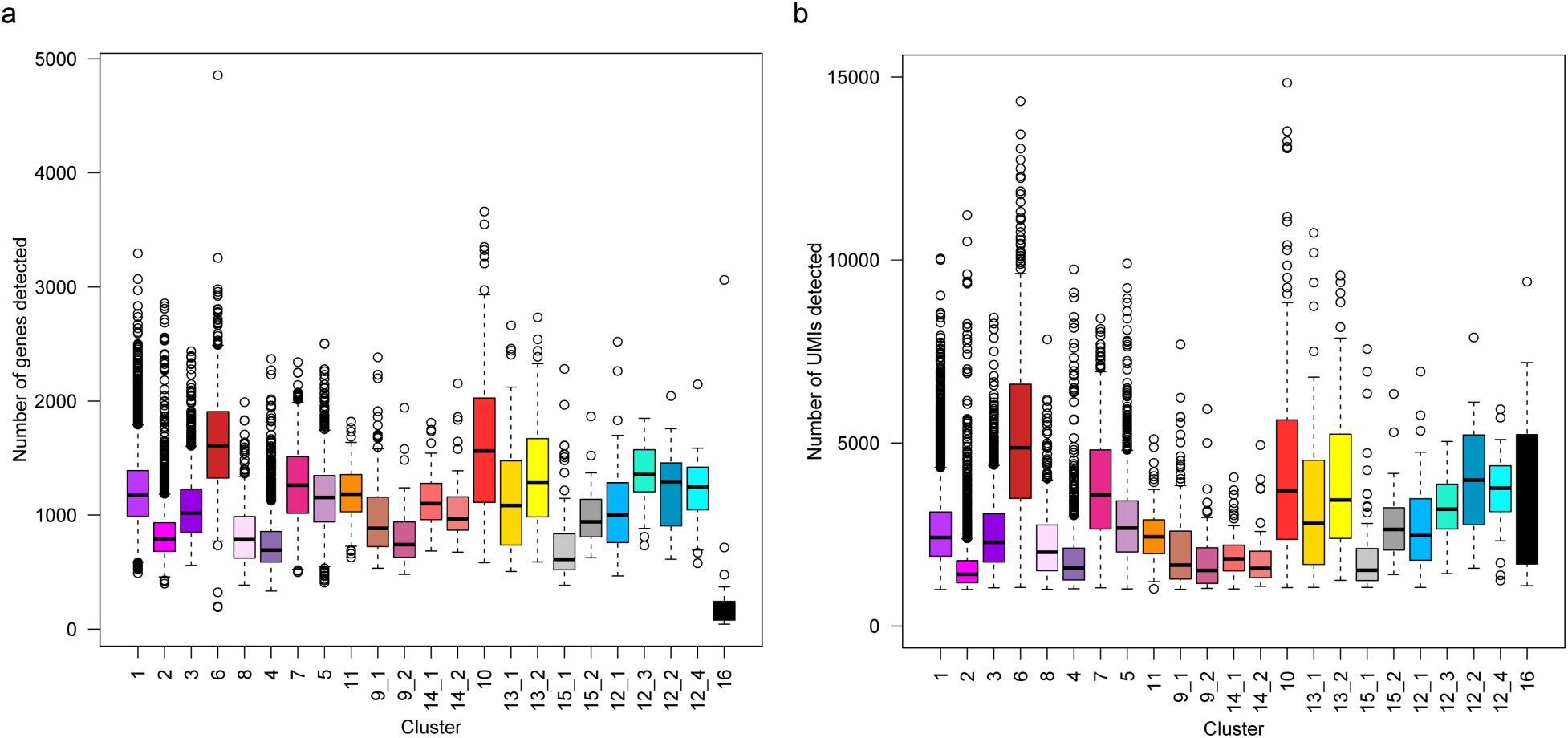
Comparability of the different clusters. **a-b** The identified clusters were comparable in terms of number of detected genes (**a**) as well as the number of detected UMIs (**b**). Abbreviations: UMI unique molecular identifier.

**Supplementary figure 2.**
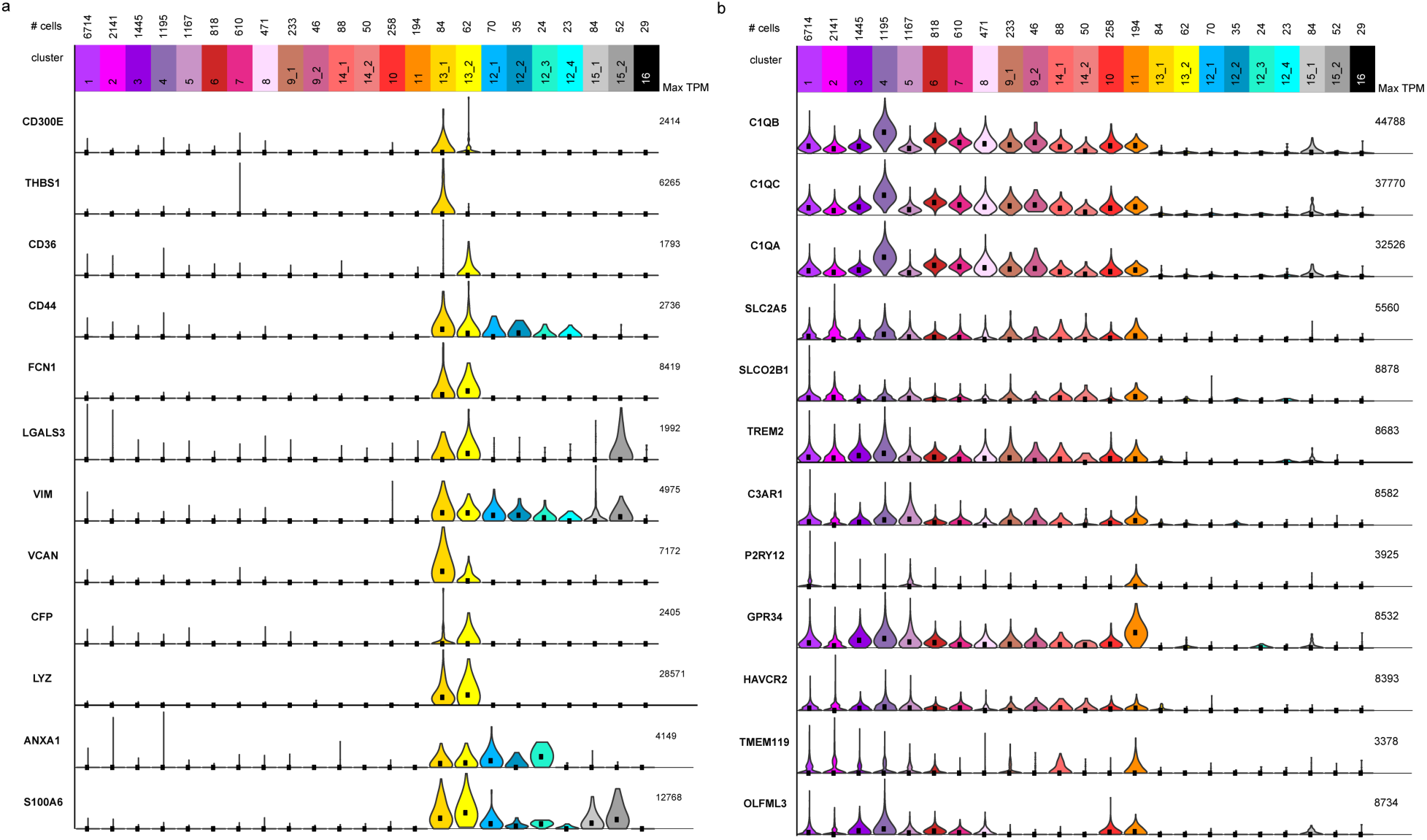
Identity of the myeloid clusters. **a** Violin plots representing the gene expression of genes that have been shown to be monocyte specific, when compared to microglia in a human study^4^. **b** Violin plots depicting the expression of genes that were found to be enriched in microglia when compared to monocytes in earlier mouse and human studies^4^,^25^,^26^,^27^. Abbreviations: TPM Transcripts Per Million; # cells number of cells.

**Supplementary figure 3.**
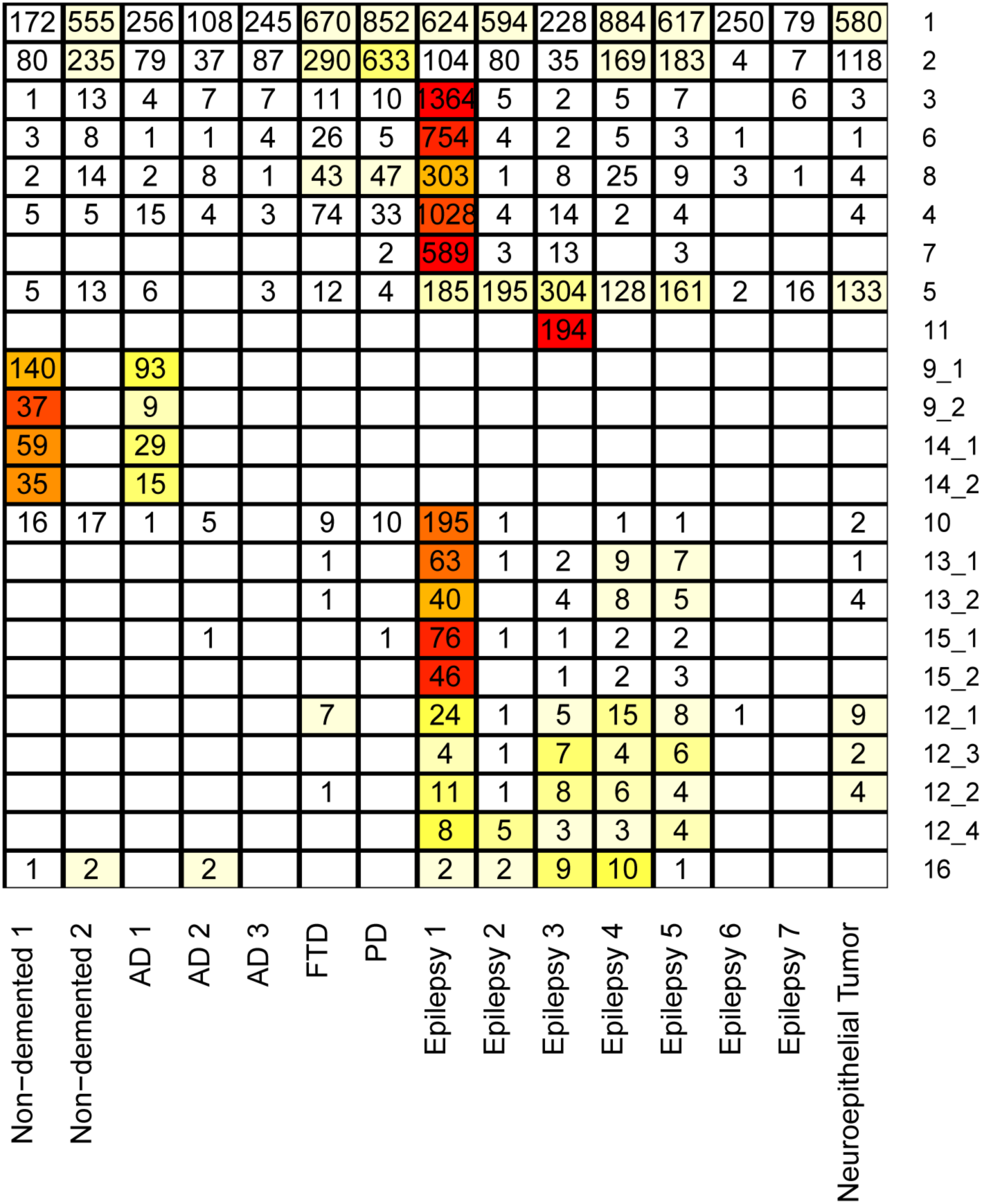
Cellular makeup of the clusters by donors. Heatmap depicting (row-wise) the number of cells each donor contributed to the given cluster; (column-wise) the cluster composition of the cells originating from each donor. Please note that clusters 1, 2, 3, 6, 8, 4 and 5 were present in all donors, cluster groups 9 and 14 were only present in some of the samples originating from aged individuals, while cluster groups 12 and 13 were only present in the biopsy samples.

**Supplementary figure 4.**
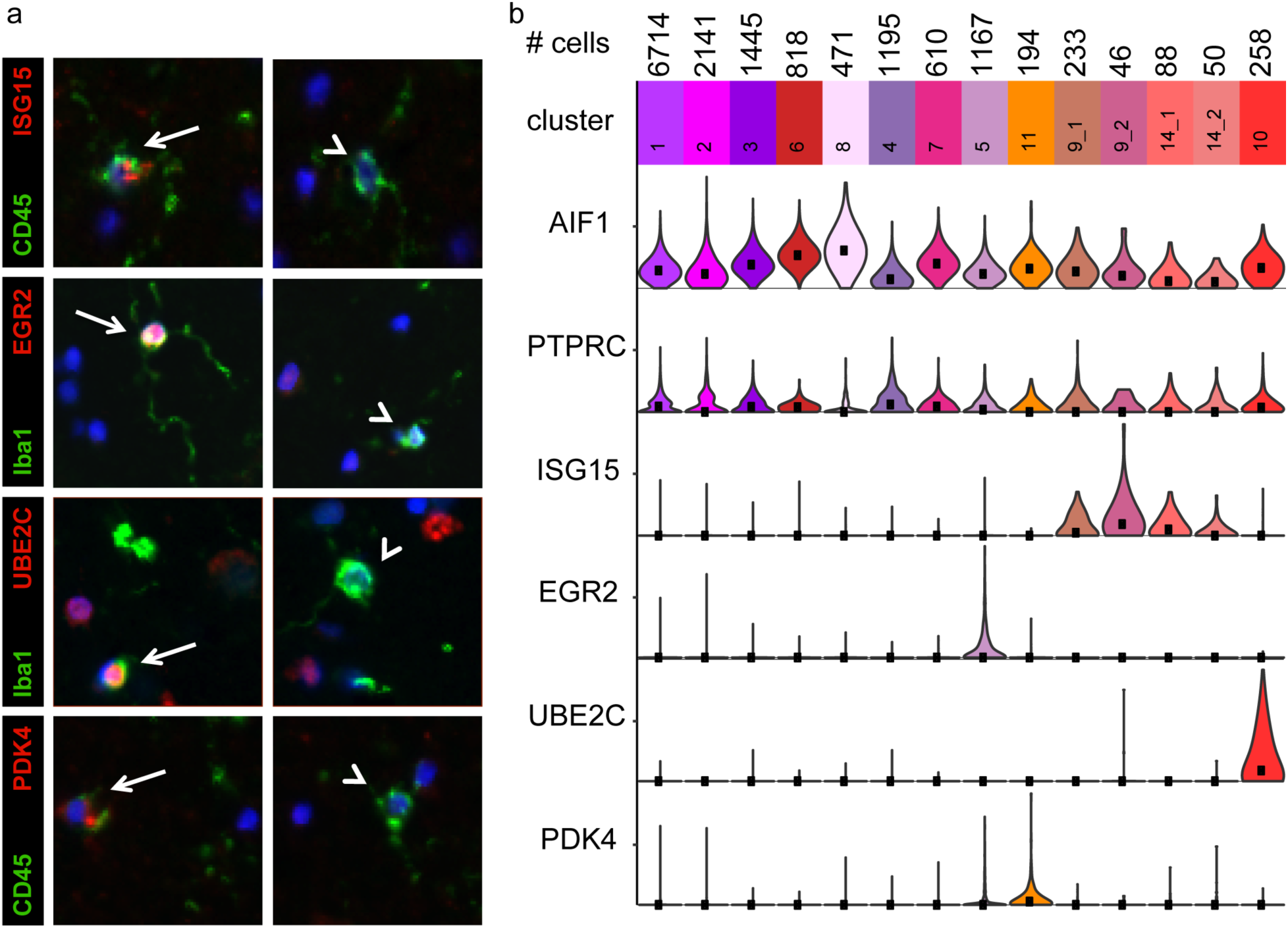
In situ confirmation of the major microglial subsets. **a** Immunofluorescence analysis confirmed that in the human brain the markers unique to 5, 11, 10 and the cluster groups 9 & 14 marked subsets of microglia (identified by either Iba1 or CD45 staining) respectively. **b** Violin plot depicting the expression of the selected markers in the different microglial clusters. AIF1 is the gene encoding the protein AIF1, also called IBA1. PTPRC is the gene encoding the surface marker CD45.

**Supplementary figure 5.**
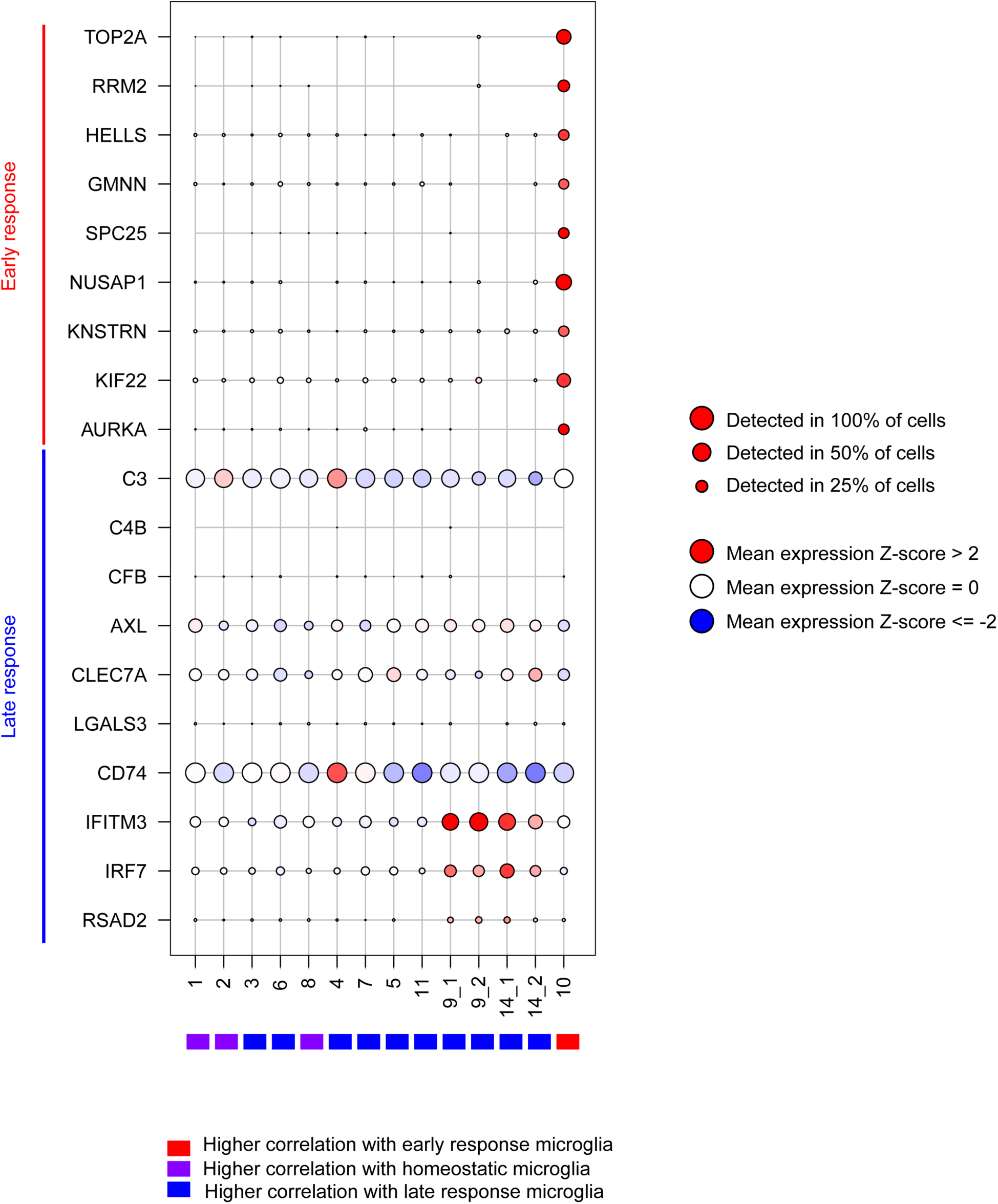
Cluster annotation based on previously published mouse studies – part 1. Figure depicting the expression levels of genes related to the early and late response mouse microglia phenotypes described in Mathys et al. 2017. The size of the dots is proportional to the number of cells expressing the given gene in the corresponding cluster. The color of the dots represents the mean Z score of expression. Bars on the bottom indicate whether the mean cluster expression correlated more strongly with the different murine microglia RNA-seq expression profiles (from Mathys et al. 2017).

**Supplementary figure 6.**
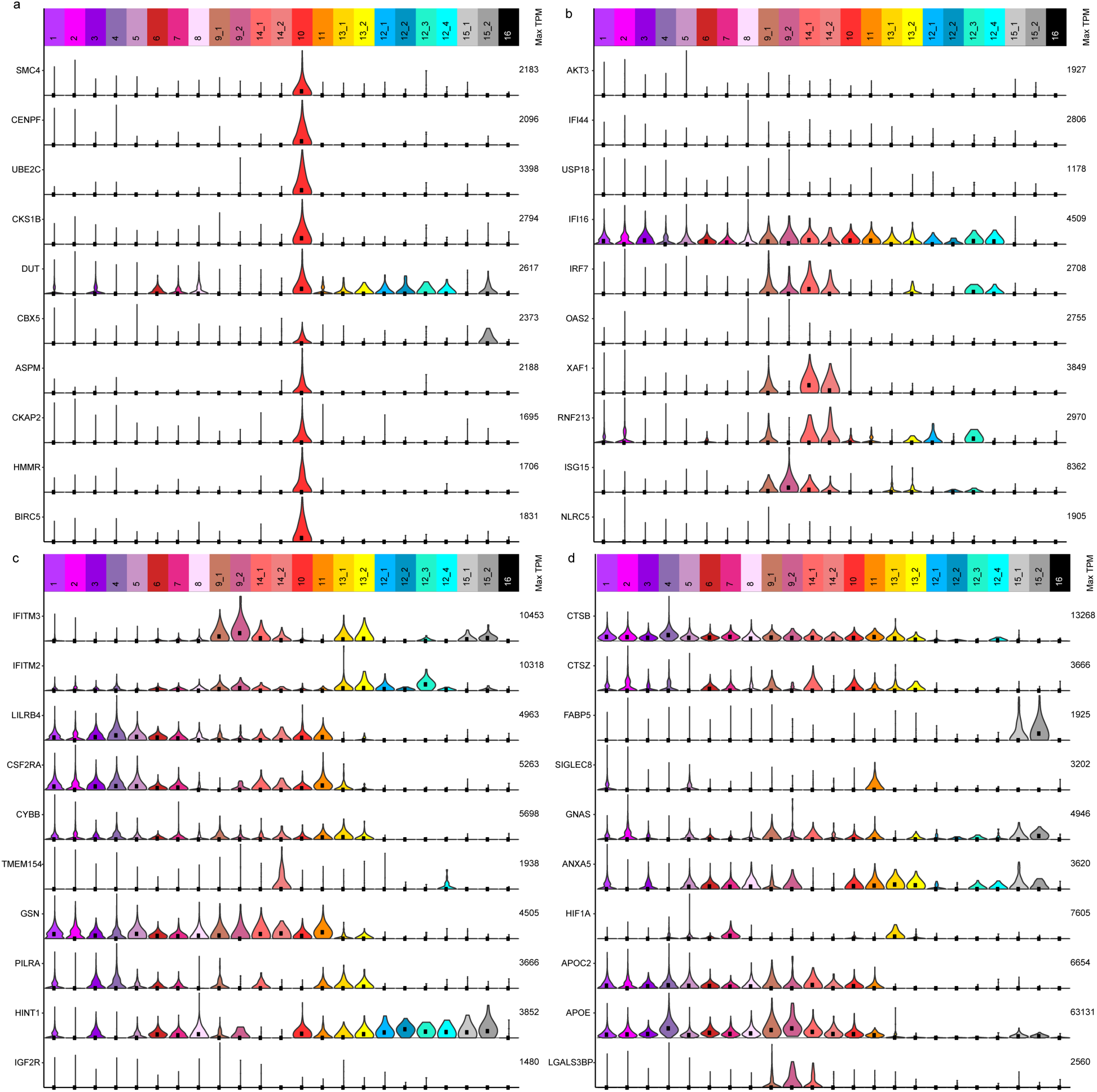
Cluster annotation based on previously published mouse studies – part 2. **a** Violin plots visualizing the genes associated with the proliferation related gene co-expression module in Friedman et al. 2018. **b** Genes associated with the Interferon related gene co-expression module from the same study. **c** Violin plot showing the signature genes of the gene co-expression module associated with LPS response (Friedman et al. 2018). **d** Neurodegeneration related genes from Friedman et al. 2018. Please note that apart from the proliferation related gene sets from both studies (**Supplementary figure 5a and Supplementary figure 6a**) the other gene sets identified in mouse fail to highlight a single human microglia subset. Abbreviations: TPM Transcripts Per Million.

**Supplementary figure 6. Cluster and donor wise expression of the DAM signature in human microglia subsets. a** Violin plots depicting the expression pattern of the homeostatic and DAM signature genes in the different human microglia subsets. **b** Violin plots showing the expression pattern of the homeostatic and DAM signature genes in the microglia of different donors that participated in our study. Please not the general expression of the DAM signature genes across the different human microglia subsets (**a**) and across the different donors (**b**). Abbreviations: TPM Transcripts Per Million.

**Supplementary figure 7.**
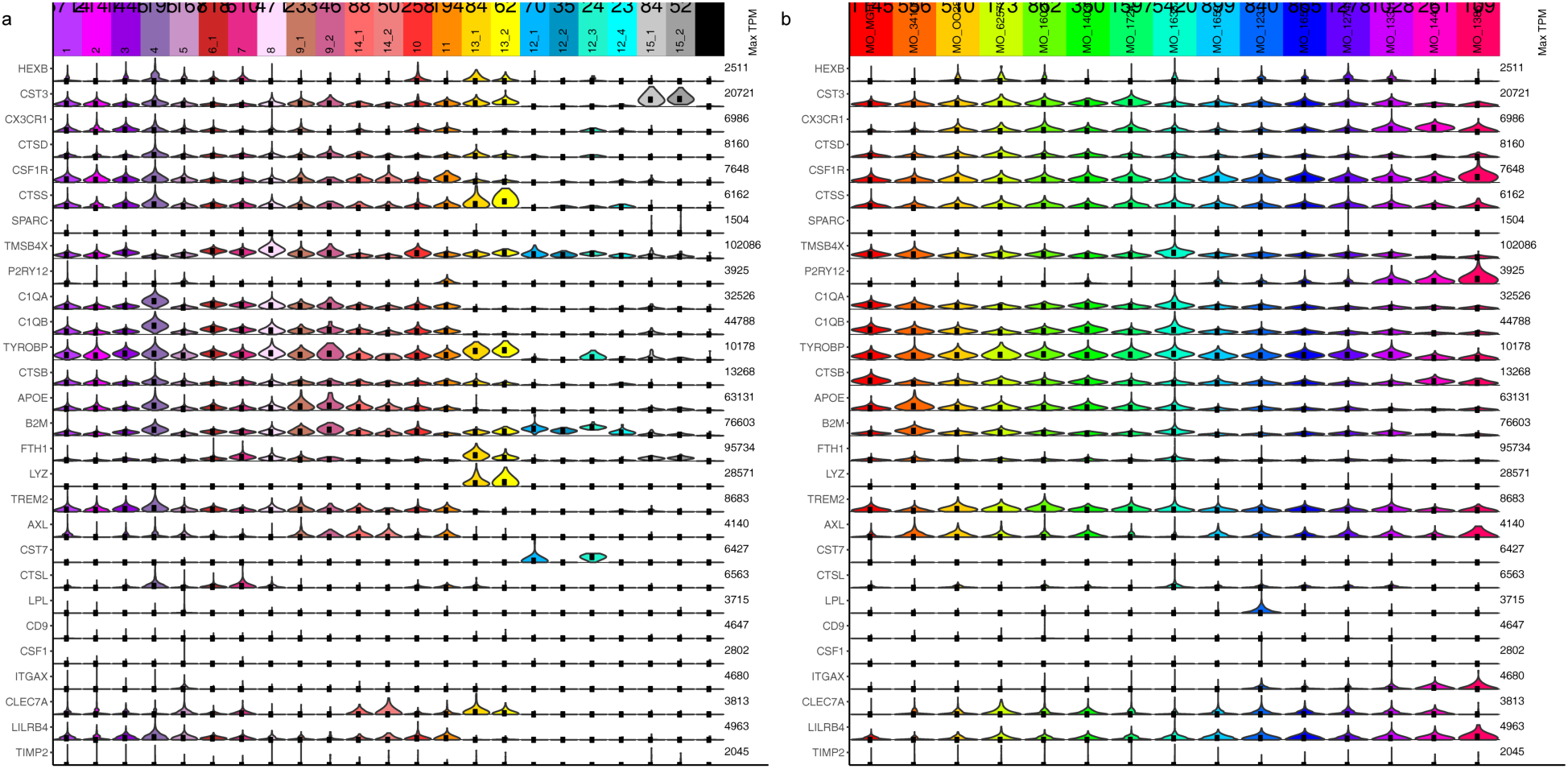
Expression of the mouse DAM signature genes in the human microglia subsets and donors. Violin plots depicting the expression distribution of the different homeostatic genes and DAM genes in the human microglia clusters (**a**) and across the different donors (**b**). Please note that the genes that were detected in our dataset were present in all the clusters and all the donors. Abbreviations: TPM Transcripts Per Million.

**Supplementary figure 8.**
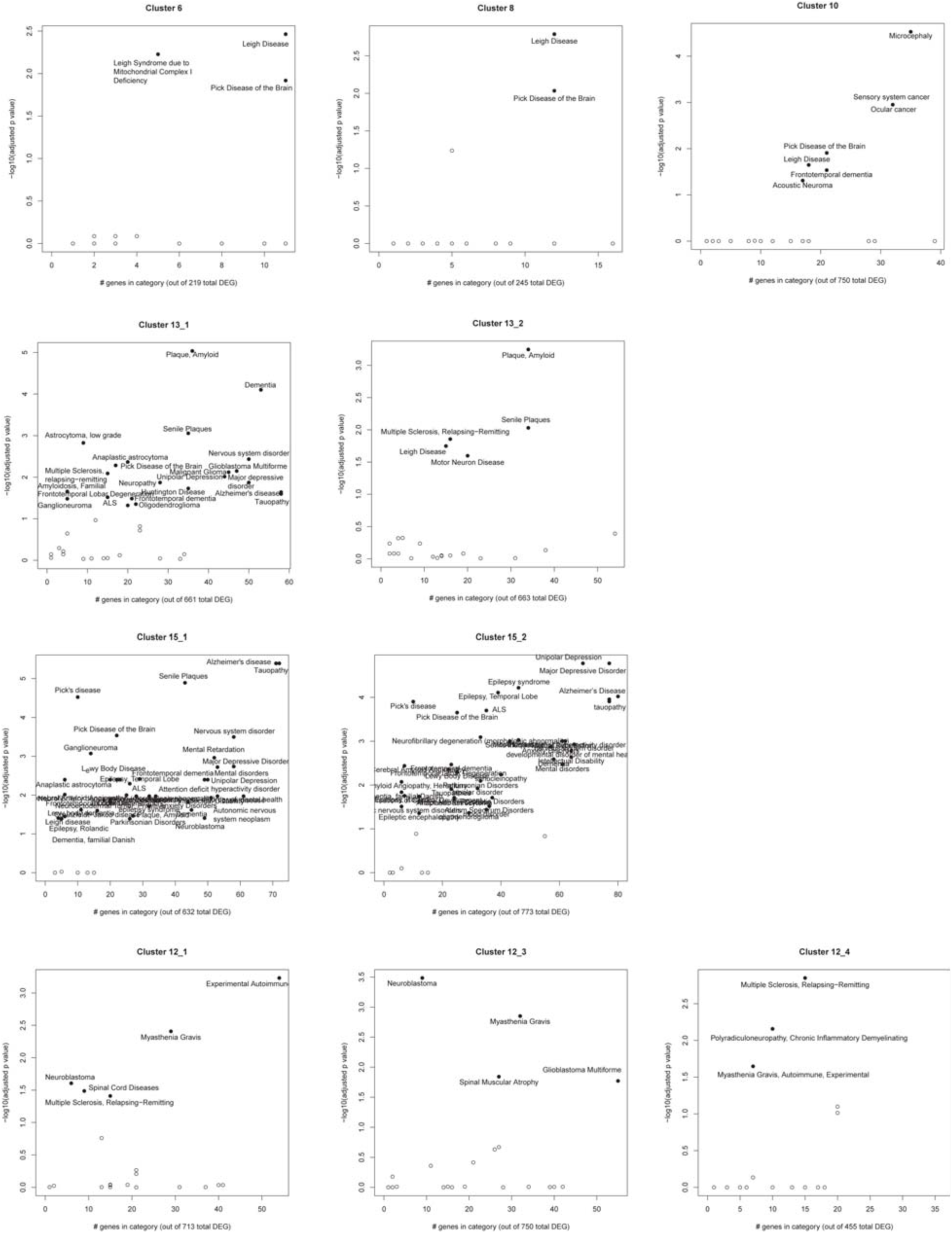
Disease association of the microglial and the non-microglial subsets. Scatter plots depicting brain related diseases that are significantly associated with the different clusters based on an enrichment analysis between disease associated gene sets from the disease ontology database (http://disease-ontology.org/) and the signature gene sets of each microglia cluster.

**Supplementary figure 9.**
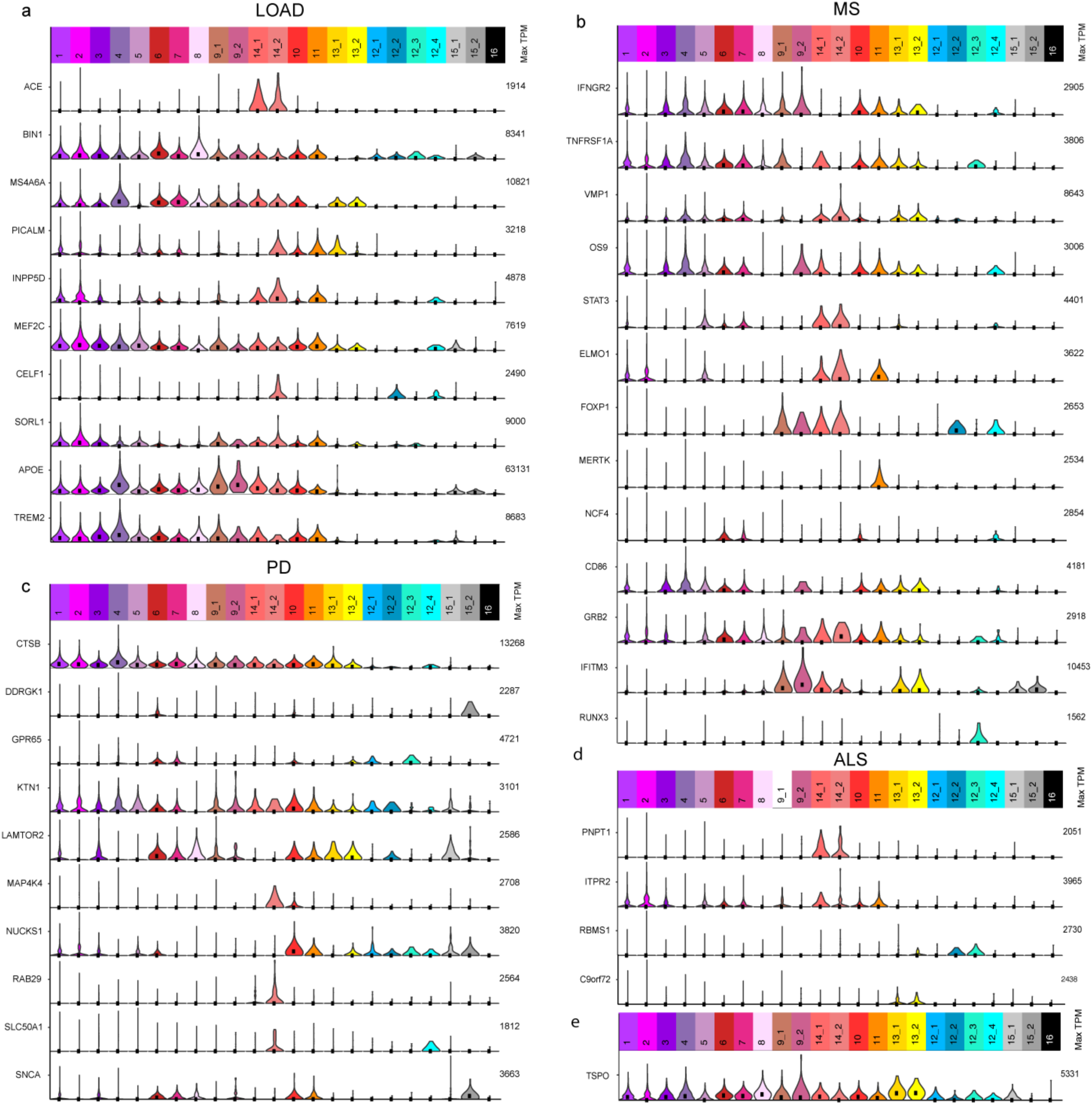
The expression of neurodegenerative disease risk genes in the different clusters. Violin plots showing the distribution of expression levels of genes associated with LOAD (**a**), MS (**b**) and PD (**c**) and ALS (**d**) (from the GWAS catalog (https://www.ebi.ac.uk/gwas/)), respectively. Only those genes are shown that had a detectable level of expression in at least one of the clusters. Abbreviations: TPM Transcripts Per Million; LOAD late onset Alzheimer’s disease; MS multiple sclerosis; PD Parkinson’s disease; ALS amyotrophic lateral sclerosis.

**Supplementary figure 10.**
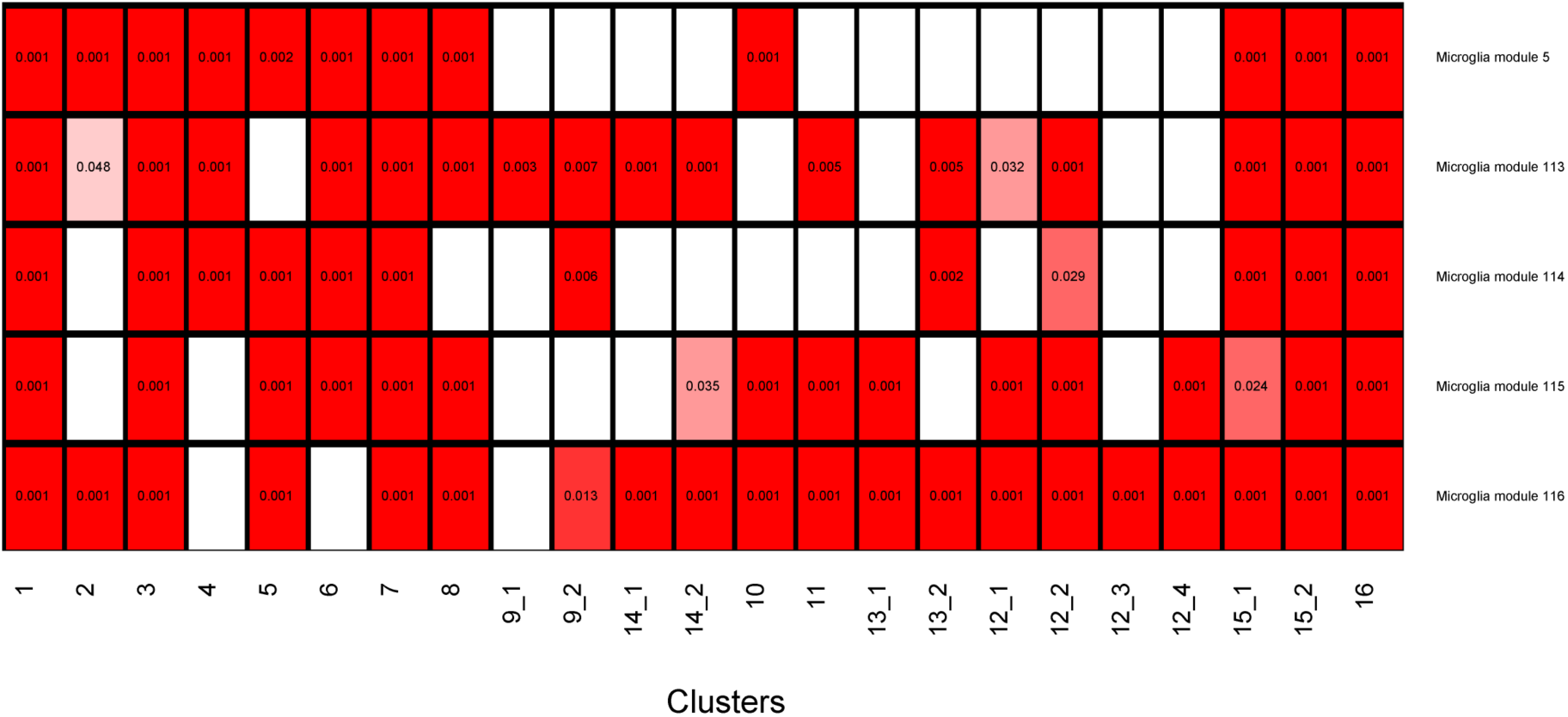
Association of microglia subsets with gene co-expression modules of the aging human brain. Heatmaps showing the association between the microglia clusters and the microglia related gene co-expression modules described in detail in Ellis et al. The numbers in the boxes represent the adjusted p-value from standard gene-set enrichment analysis.

## Methods

### Source of human brain specimens

Five autopsy brain specimens originated from the Rush University Medical Center/Rush Alzheimer’s Disease Center (RADC) in Chicago, IL from our collaborator Dr. Bennett. The one Parkinson’s disease autopsy sample originated from Banner Sun Health Research Institute in Phoenix, AZ through our collaborator Dr. Beach. The one frontotemporal dementia autopsy sample was received from Drs. Hyman and Frosch from Massachusetts General Hospital, Boston MA. The seven surgically resected brain tissue specimens originated from the Brigham and Women’s Hospital in Boston, MA fromcollaborators Drs. Sarkis, Cosgrove, Helgager, Golden, and Pennell. All brain specimens were obtained through informed consent and/or brain donation program at the respective organizations. All procedures and research protocols were approved by the corresponding ethical committees of our collaborator’s institutions as well as the Institutional Review Board (IRB) of Columbia University Medical Center. For a detailed description of the brain regions sampled, age of the donors, histopathology and clinical diagnosis please see **Supplementary data 1**.

### The ROS and MAP cohorts at RADC

Five out of seven autopsy specimens used in this study originated from two prospective studies of aging: the Religious Orders Study (ROS)^1^ and the Memory and Aging Project (MAP)^2^. Participants to enter these prospective studies have to be at least 53 (ROS) or 55 (MAP) years old and non-demented at the time of enrollment and sign an Anatomical Gift Act agreeing to donate their brain and spinal cord at the time of death. Each subject undergoes annual neuropsychologic evaluations while alive and a structured, quantitative neuropathologic examination at autopsy. Brain specimens were distributed for this project from autopsies taking place Sunday morning to Thursday. Only autopsies for which the post mortem delay was less than 12 hours were included in this study.

### Shipping of brain specimens

After weighing, the tissue was placed in ice-cold transportation medium (Hibernate-A medium (Gibco, A1247501) containing 1% B27 serum-free supplement (Gibco, 17504044) and 1% GlutaMax (Gibco, 35050061)) and shipped overnight at 4°C with priority shipping.

### Microglia isolation and sorting

The isolation of microglia was performed as described elsewhere ^3^. Breifly, upon arrival of the autopsy sample, the cortex and the underlying white matter were dissected under a stereomicroscope. The epilepsy surgery samples were processed without dissection as in this case the cortical white and grey matter were not always distinguishable. All procedures were performed on ice. From the autopsy sample only microglia isolated from the grey matter were used in this study. The dissected tissue was placed in HBSS (Lonza, 10-508F) and weighed. Subsequently the tissue was homogenized in a 15 ml glass tissue grinder - 0.5g at a time. The resulting homogenate was filtered through a 70 um filter and spun down with 300g for 10 minutes. The pellet was resuspended in 2 ml staining buffer (PBS (Lonza,) containing 1% FBS) per 0.5g of initial tissue and incubated with anti-myelin magnetic beads (Miltenyi, 130-096-733) for 15 minutes according to the manufacturers specification. The homogenate was than washed once with staining buffer and the myelin was depleted using Miltenyi large separation columns. The cell suspension was spun down and incubated with anti-CD11b magnetic beads (Miltenyi, 130-049-601) for 15 minutes. Following a wash with staining buffer the CD11b+ cells were isolated on a Miltenyi MS column while the CD11- fraction was cryopreserved using FBS containing 10% DMSO. The CD11b positive fraction was than incubated with anti-CD11b AlexaFluor488 (BioLegend, 301318) and anti-CD45 AlexaFluor647 (BioLegend, 304018) antibodies as well as 7AAD (BD Pharmingen, 559925) for 20 minutes on ice. Subsequently the cell suspension is washed twice with staining buffer, filtered through a 70 um filter and the CD11b+/CD45+/7AAD- cells (Supplementary figure 1b) were sorted on a BD FACS Aria II sorter. Cells were sorted in A1 well of a 96 well PCR plate (Eppendorf, 951020401) containing 100 ul of PBS buffer and submitted to single cell library construction.

### 10x Genomics Chromium single cell 3’ library construction

Viability was assessed by trypan blue exclusion assay and cell density was adjusted to 175 cells per µl. 7,000 cells were then loaded on a single channel on the 10x Chromium chip from each sample. The 10x Genomics Chromium technology enables 3’ digital gene expression profiling of thousands of cells from a single sample by separately indexing each cells transcriptome. First, thousands of cells will be partitioned into nanoliter-scale Gel Bead-In-EMulsions (GEMs). Within one GEM all generated cDNA share a common 10x barcode. Libraries are generated and sequenced from the cDNA and the 10x barcodes are used to associate individual reads back to the individual partitions. To achieve single cell resolution, the cells are delivered at a limiting dilution. Upon dissolution of the Single Cell 3’ Gel Bead in a GEM, primers containing (i) an Illumina R1 sequence (read 1 sequencing primer), (ii) a 16 nt 10x Barcode, (iii) a 10 nt Unique Molecular Identifier (UMI), and (iv) a poly-dT primer sequence are released and mixed with cell lysate and Master Mix. Incubation of the GEMs then produces barcoded, full-length cDNA from poly-adenylated mRNA. After incubation, the GEMs are broken and the pooled fractions are recovered. Full-length, barcoded cDNA is then amplified by PCR to generate sufficient mass for library construction. Enzymatic fragmentation and size selection are used to optimize the cDNA amplicon size prior to library construction. R1 (read 1 primer sequence) are added to the molecules during GEM incubation. P5, P7, a sample index, and R2 (read 2 primer sequence) are added during library construction via end repair, A-tailing, adaptor ligation, and PCR. The final libraries contain the P5 and P7 primers used in Illumina bridge amplification. The described protocol produces Illumina-ready sequencing libraries. A Single Cell 3’ Library comprises standard Illumina paired-end constructs which begin and end with P5 and P7. The Single Cell 3’ 16 bp 10x Barcode and 10 bp UMI are encoded in Read 1, while Read 2 is used to sequence the cDNA fragment. Sample index sequences are incorporated as the i7 index read. Read 1 and Read 2 are standard Illumina sequencing primer sites used in paired-end sequencing. Sequencing the generated library produces a standard Illumina BCL data output folder. The BCL data will include the paired-end Read 1 (containing the 16 bp 10x Barcode and 10 bp UMI) and Read 2 and the sample index in the i7 index read.

### Batch structure and sequencing

The fresh autopsy and surgical resection samples were processed for microglia isolation, library construction and sequencing as they were obtained. Accordingly each sample constitute one batch for all three procedures. All sequencing was performed on an Illumina HiSeq4000 machine. For specifics regarding the generated reads see **Supplementary data 2**.

### Computational methods

Barcoded reads were demultiplexed and aligned to the GRCh38 genome with Ensemble transcriptome annotation (downloaded June 2017, GRCh38.85) using CellRanger with default parameters. Only cells with >1000 Unique Molecular Identifiers (UMIs) were kept for clustering and downstream analysis. Putative cell types were identified using an iterative clustering approach; after regressing out batch and total UMI number, all genes with variance greater than the mean were used to cluster cells with the PCA-Louvain clustering approach, as implemented in the Seurat R package^4^. After clustering, cluster robustness was assessed by training a classifier on half the cells and predicting the cluster membership of the remaining half. Any clusters that showed a minimum less than 50% accurate prediction over 20 iterations were merged. Each cluster was then iteratively subclustered using the same procedure until no further robust clusters were found. After clustering, differential genes were identified over all pairs of clusters using the edgeR package^5^.

Constellation diagrams showing the relationship among different clusters were generated using a cross-validation machine learning approach^6^. For each pair of clusters, cells were classified using five-fold cross-validation using a random forest classifier (trained on 80% of the cells). This process was repeated 10 times, resulting in a membership score for each cell belonging to one or the other cluster in the pair. Cells that were not unambiguously classified (10 times out of 10) to the same cluster were called “intermediate” cells. For the constellation diagram, the edges between any two clusters represent the percentage of total cells (from the pair of clusters) that were called “intermediate”, and the size of the nodes represents the total (core+intermediate) cells originally assigned to that cluster.

For dotplot representations, expression values for a given gene were z-scored over all the cells belonging to all the clusters visualized, and then per-cluster means of the z-scored values were calculated and plotted using the color scheme shown in each figure. Sizes of the circles represent the number of cells in the cluster in which the gene was detected (TPM>0).

For all gene-based association tests, we first obtained lists of cluster-specific genes; for each cluster, this comprised all genes that showed statistically significant up-regulation in comparison to at least one cluster, with the constraint that the gene did not show significant down-regulation with respect to any other cluster. Using these cluster-specific gene lists, we performed gene ontology analysis using the topGO R package^7^ with standard parameters and Benjamini-Hochberg procedure to control the FDR rate. Disease ontology analyses were performed using the DOSE R package^8^, also with standard parameters and Benjamini-Hochberg procedure. To assess enrichment cluster-specific signatures in previously-reported microglial modules derived from bulk data^9^,^10^. Gene-set enrichment analyses were performed with Mann-Whitney test to assess whether the module-specific genes showed a different distribution of minimum p-values, as compared to non-module-specific genes. Significance of Mann-Whitney p-values were compared to 10000 random shufflings of the cluster-specific gene p-values, and then Bonferroni-corrected to obtain adjusted p-values. For the association of cluster-specific genes with the ROSMAP trait-associated genes, we ran rank-rank hypergeometric test^11^, using increments of 20 genes in each list (cluster-specific genes and trait-associated genes), finding the minimum p-value for significant overlap. These p-values were then compared to the minimum p-values for significant overlap over 10000 random shufflings of the two gene sets, and Bonferroni-corrected to obtain adjusted p-values.

### Immunohistochemistry

Immunohistochemistry was performed as described elsewhere^10^. Briefly, 6 µm thick sections of human pre-frontal cortex were de-paraffinized with Xylene for 20 minutes. The sections were put through an ethanol series (ethanol 100%, ethanol 100%, ethanol 70% - 1 minute for each) and re-hydrated in water (for 1 minute). Subsequently, the slides were washed 3 times with phosphate buffered saline (PBS). Antigen retrieval was achieved by putting slides in pH 6.0 citrate buffer and using microwave for 25 min at 400 Watt. The slides were placed in tap water for 5 minutes, washed three times with PBS. Unspecific binding of antibodies was blocked with 3% bovine serum albumin (BSA) in PBS containing 0.1% TritonX for 20 min. Primary antibody was applied overnight. Subsequently the slides were washed with PBS three times and the fluorochrome conjugated secondary antibody was applied to the slides for one hour. The slides were again washed three times with PBS. Endogenous autofluorescence was quenched with sudan black for 10 minutes. The slides were again washed with BPS three times and mounted with ProlongGold containing DAPI. The primary antibodies used were rabbit anti-human Iba1 (Wako; 019-19741; at the dilution of 1:500), mouse anti-human CD45 (Novus; NB500-319; 1:200), rabbit anti-human ISG15 (Proteintech; 15981-1-AP; 1:100), mouse anti-human EGR2 (OriGene; TA505382; 1:50), mouse anti-human UBE2C (Proteintech; 66087-1-IG; 1:50), rabbit anti-human PDK4 (Proteintech; 12949-1-AP; 1:50). The secondary antibodies used were goat anti-mouse IgG (H+L) highly cross-adsorbed secondary antibody conjugated to Alexa Fluor Plus 488 (ThermoFisher Scientific; A32723; 1:300) or Alexa Fluor Plus 555 (ThermoFisher Scientific; A32727; 1:300) and goat anti-rabbit IgG (H+L) highly cross-adsorbed secondary antibody conjugated to Alexa Fluor Plus 488 (ThermoFisher Scientific; A32731; 1:300) or Alexa Fluor Plus 555 (ThermoFisher Scientific; A32732; 1:300).

